# Bioarchaeological evidence for commensal brown rats (*Rattus norvegicus*) in early Neolithic China

**DOI:** 10.64898/2026.01.03.697376

**Authors:** Thomas Cucchi, Ardern Hulme-Beaman, David Orton, Peng Lyu, Li Zhipeng, Yuan Jing

## Abstract

Due to its commensal relationship with humans, facilitating its dispersal from its natural range in East Asia, the brown rat (*Rattus norvegicus*) is now one of the most cosmopolitan and invasive mammals. Brown rats are also vectors of human pathogens and the most commonly used mammalian model (after mice) for biomedical research. Despite their major role in biodiversity loss and medicine, our understanding of a likely origin for their commensal relationship with humans in China still relies on modern population genetics, without a reliable timeframe of localisation. In this research, we have tried to fill this gap using direct information from the archaeological remains of rats collected by Chinese colleagues in several archaeological sites dated between 9000 and 3500 BP. Thanks to high-resolution morphometrics and direct ^14^C dating, we have identified the occurrence of the brown rat (*Rattus norvegicus)* within the domestic area deposits of the early Neolithic site of Jiahu by 7500 cal BP, suggesting a tentative link between rice agriculture and an early commensal behaviour of this species.

## Introduction

The brown rat (*Rattus norvegicus*) is currently present in almost every human environment across the globe, including every continent bar Antarctica (Musser & Carleton 2005). Yet this species’ natural range was restricted to Eastern Asia, until some populations of brown rat engaged in a commensal trajectory with humans. Benefiting from human trade and exchange networks across land and seas, brown rats were able to colonize every corner of the globe and become one of the most invasive and threatening species for biodiversity (Capizzi et al. 2014). Both economically and ecologically damaging, the species has long been associated with damage to agricultural crops, stored foodstuffs of all descriptions (Stenseth et al. 2003) and also predation of native flora and fauna (Veitch & Clout, 2002). A notable nest predator, its arrival in new environments, particularly islands, has resulted in the dramatic decline of resident bird populations (Jones et al. 2008; Sarmento et al. 2014) making the brown rat a major conservation concern and subject to extensive extirpation efforts (Calmet et al. 2001, Russel & Holmes, 2015). It is also a major source of zoonotic diseases, harbouring and spreading some of the most dangerous and infectious pathogens that afflict human societies around the world (Webster et al. 1995; Sunbul et al. 2001; Heyman et al. 2011).

Brown rats are not just an ecological plague for global biodiversity. They also provide many services for humanity. They are currently one of the main animal models for experimental research in embryology, bacteriology, virology, oncology, pharmacology and geriatrics. Populations of laboratory brown rat have facilitated many major advances in medical knowledge and treatment by the development, through inbreeding, of phenotypes similar to human diseases (Suckow et al. 2005). More recently, they became a model in evolutionary research to better understand the effect of urbanization in species evolution (Munshi-South et al. 2016). Less known is the service they provide as commensal populations occupying the sewage systems of the major cities worldwide; in Paris for example, brown rats are estimated to consume 850 tonnes of refuse per year and clean the sewage system (unpublished data).

Despite this significant impact on ecosystems, national economies and human health, the history of its early commensal interaction with human societies and dispersal is poorly known (Hulme-Beaman et al. 2021, Munshi-South et al. 2024). Thanks to bioarchaeological and historical research, we have a fairly clear picture for the dispersal timing of the brown rats in Europe and North America (Fig 1). Yet we are still lacking bioarchaeological evidence for the emergence of its commensalism and dispersal with humans in East Asia. Genetic and palaeontological studies suggest that *Rattus norvegicus* is native from Northern China and Mongolia, originating sometimes during the Pleistocene (Zeng et al. 2014, Puckett & Munshi-South 2019, Smith & Xie 2008), with fossil occurrences in upper Pleistocene and Holocene caves’ natural deposits in Sichuan-Guizhou (Zheng 1993). Genomics studies suggest that human-mediated dispersal of brown rats across South East Asia from this natural range occurred between the 9^th^ and the 16^th^ century, with dispersal towards Europe via the Middle East potentially starting through maritime trade from the 13^th^ century AD (Puckett & Munshi-South 2019). There are putative archaeological occurrences of Brown rat in 14^th^ c. AD Italy (Clark 1989) and several medieval sites in northern Europe (Ervynck 1989) that may suggest the presence of small temporary pioneer populations in Europe around that time but historical and archaeological sources rather support an invasive dispersal in Western Europe around the 18^th^ c. AD. Historical sources relating to Dublin (1729, reproduced in Wardell 1901; 1722 according to Rutty, 1772), Bornholm (Urne 1755, cited in Winge 1908), London (Smith 1768), Paris (Buffon, 1760), western Norway^1^ (1762, Strøm n.d. cited in Collett 1912) and Braunschweig (Zimmerman 1777), all note the arrival to Europe of a new species of rat that was both larger and displayed more aggressive behaviour compared to the black rat (*R. rattus*), present in Europe over the preceding ∼2,000 years (Armitage 1994; Vigne & Valladas 1996; Audoin-Rouzeau & Vigne 1997). The arrival of the brown rat led to the widespread replacement of the black rat in Europe, owing in part to the brown rat’s greater tolerance of colder climates (Armitage 1994) and greater competitive aggression considering recent studies (Himsworth et al. 2014; Feng & Himsworth 2014). These historical accounts fit well with the earliest firm archaeological evidence for the arrival of brown rat in Europe, a skull taxonomically identified from a 1796 AD shipwreck off the Mediterranean Coast of Corsica Island (Vigne & Villié 1995).

**Figure 1.**
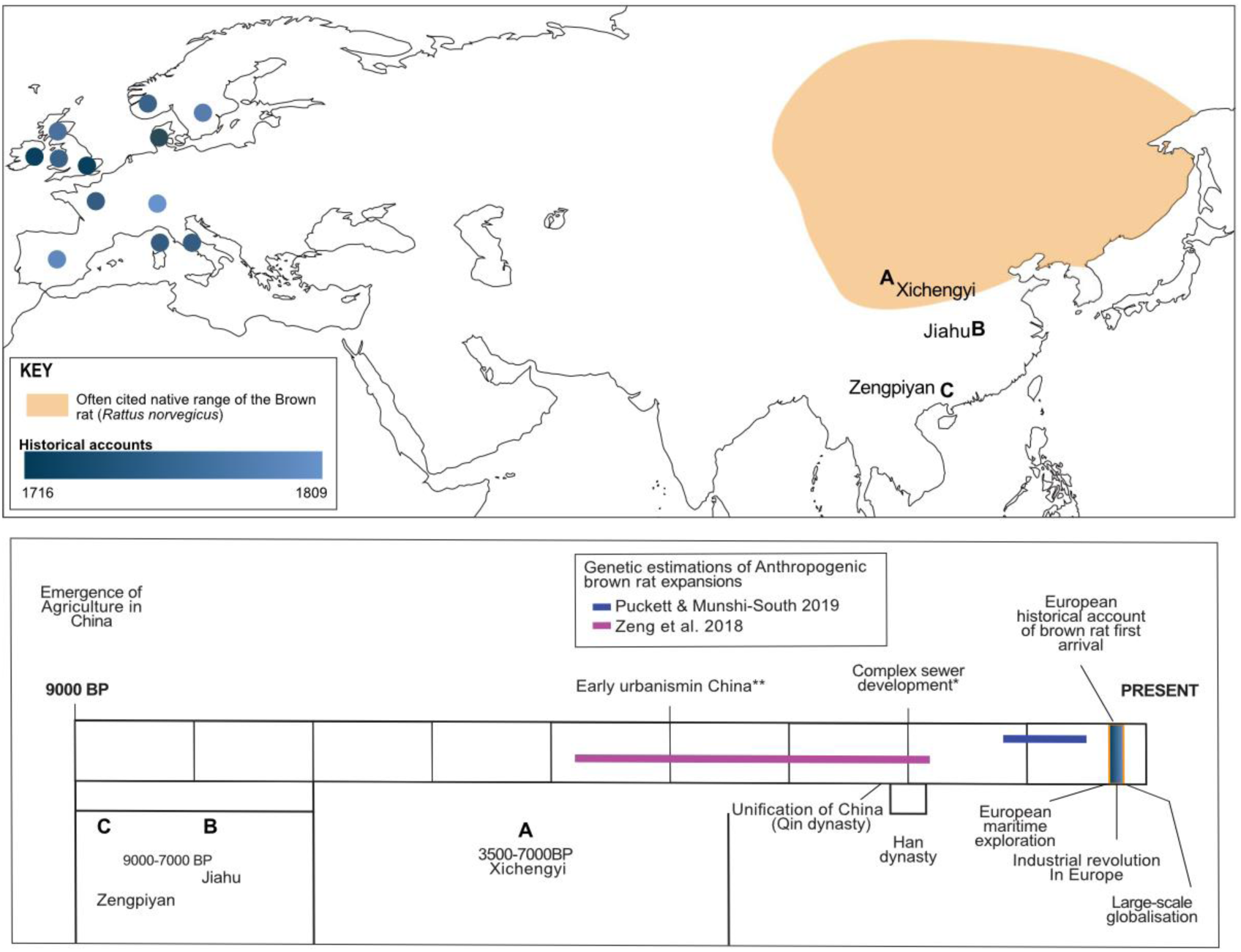
Timeline and mapping of historical and archaeological evidence for commensal brown rats in Europe and locations of the Chinese archaeological sites examined here: A) Xichengyi (Yanbo et al. 2014); B) Jiahu (Juzhong, Z. & Qilong C. 2013); C) Zengpiyan (Zhang 1990). Distribution of native brown rat in orange after Hedrich (2006).

From Europe, brown rats appear to have spread rapidly to European colonies in North America. Securely dated finds are reported from a 1760 shipwreck in New Brunswick, along with frequent confirmed identifications at terrestrial sites on the Atlantic coast from at least the mid-18^th^ c. (Guiry et al. 2024). At the same time there is evidence for a separate expansion to Pacific North America associated with Russian colonial activity. ‘Large grey rats’ were already reported on the Aleutian Rat Islands – later known for their *Rattus norvegicus* infestation – in the late 1770s and linked to a ‘foreign’ shipwreck (Pallas 1782), later described as Japanese (Khlebnikov, 1826-27 [1979]). A mid-1820s account from the Russian American capital at Sitka, Alaska, suggests that abundant rats – most likely *R. norvegicus* given the climate – had reached the territory with shipping from Okhotsk and/or Guangzhou (Khlebnikov, c.1825 [trans. Dmytryshyn & Crownhart-Vaughan, 1976]). Phylogenetic data support these separate dispersal events, with a deep split between lineages suggesting discrete source populations in each case (Puckett & Munshi-South 2019).

Genetic estimations from modern samples of the brown rat suggest a commensal relationship with humans by at least 800 BP in east Asia (Puckett et al. 2016; Puckett & Munshi-South 2019) though other studies suggest a much earlier date or at least >3500 BP (Zheng 2015). However, these occurrences have not been supported by taxonomic identification with high resolution morphometric approaches or extensively tested with ancient DNA comparisons, let alone form direct ^14^C dates. Separate genomic studies have also produced somewhat differing results with some discrepancies between a likely north or south east Asian origin of Holocene *Rattus norvegicus* (Song et al. 2014; Zeng et al. 2018; Puckett & Munshi-South 2019). Given the complex history of the relationship between humans and brown rats over the last ∼500 years it is maybe unsurprising that genetic estimations applied to modern samples find conflicting signals.

As recently shown for the house mouse (Weissbrod et al. 2017; Cucchi et al. 2020), without bioarchaeological evidence one cannot appreciate the role of the human niche construction in the development of this species’ commensal behaviour and the dynamic factors underlying its biological invasion history. Such insights from the archaeological record are now possible thanks to the development of numerical taxonomy using 2D geometric morphometrics of molars occlusal view which allow accurate species identification within complexes of twin species like mice (Cucchi et al. 2020) and rats (Hulme-Beaman et al. 2018). To track the origin of brown rat commensalism in East Asia, we applied these techniques to identify dental remains of *Rattini* from three archaeological sites in both Southern and Northern China (Figure 1) across the “Neolithic” transition period (10000-3000 BP).

## Materials and methods

We studied thirteen mandibular jaws of the *Rattus* genus curated at the Chinese Academy of Social Science in Beijing that were collected during the excavation of the deposits from three Chinese archaeological sites (Figure 1): Zengpiyan cave (10000–7000 BP) located in the Guilin province in Southern China (Zhang 1990), the Early Neolithic site of Jiahu (9000–7000 BP) located in the Henan Province near the Huei river (Juzhong & Qilong 2013) and the Middle Neolithic Site of Xichengyi (7000–3500 BP) in the Hexi corridor north of the lower Yellow river valley (Yanbo et al. 2014). Zengpiyan cave provides a long sequence of occupation by late hunter-gatherers with pottery, exploiting a great diversity of wild plants and animals with more than 110 species of mammals, birds, fish and reptiles identified (Shelach-Lavi 2015) with so far, no clear evidence for a transition towards plant cultivation (Denham et al. 2018) or pig domestication (Cucchi et al. 2011, 2016). Jiahu is one of the major sites of the Early Neolithic of China where the earliest evidence for sedentary villages, rice cultivation (Zhang et al. 2012) and pig domestication (Jing & Flad, 2002; Cucchi et al. 2011) has been documented. Xichengyi is a very large settlement (35 ha) which illustrates the transition between the Late Neolithic and the Bronze Age and provided a large quantity of copper metallurgy remains (Yang et al. 2019). The farming economy of Xichengyi relied on millet pluvial agriculture as well as introduced wheat and barley and animal husbandry of pigs, cattle and sheep along with hunting.

Jiahu provided a total of 40 remains and a minimum number of eight *Rattus* sp. individuals, collected from layers discovered in three excavation units (T107, T43, T42 and T41) located in the central eastern and western part of the site extension. The layers of all these units are an integral part of the domestic function of the site comprising semi-subterranean houses, storage area, waste pits and burials that allowed documentation of daily life at Jiahu; the spatial organisation of houses, waste management and food consumption (Juzhong & Qilong 2013). A minimum number of two *Rattus* sp. individuals were found in the unit T107 with one frontal bone fragment, seven appendicular bones, 2 maxillary bones, 2 incisors and one mandibula with its m1 and m2. At least 1 *Rattus* sp. individual was found in unit T41, with 2 mandibles with m1 and 2 appendicular bones (humerus and ulna). A minimum of 3 *Rattus* sp. individuals were found in unit T42 with 3 mandibles bearing an identifiable m1 and 8 appendicular bones. Finally, a minimum of 2 *Rattus* sp. individuals were found in unit T43, with 3 mandibles and 8 appendicular bones. None of the 13 mandibles analysed show signs of digestion by predators, neither nocturnal raptors nor small carnivores, and some display a brown coloration potentially due to combustion (Figure 2A), which suggest *in situ* accumulation within the settlement’s deposits rather than accumulation by predators.

**Figure 2:**
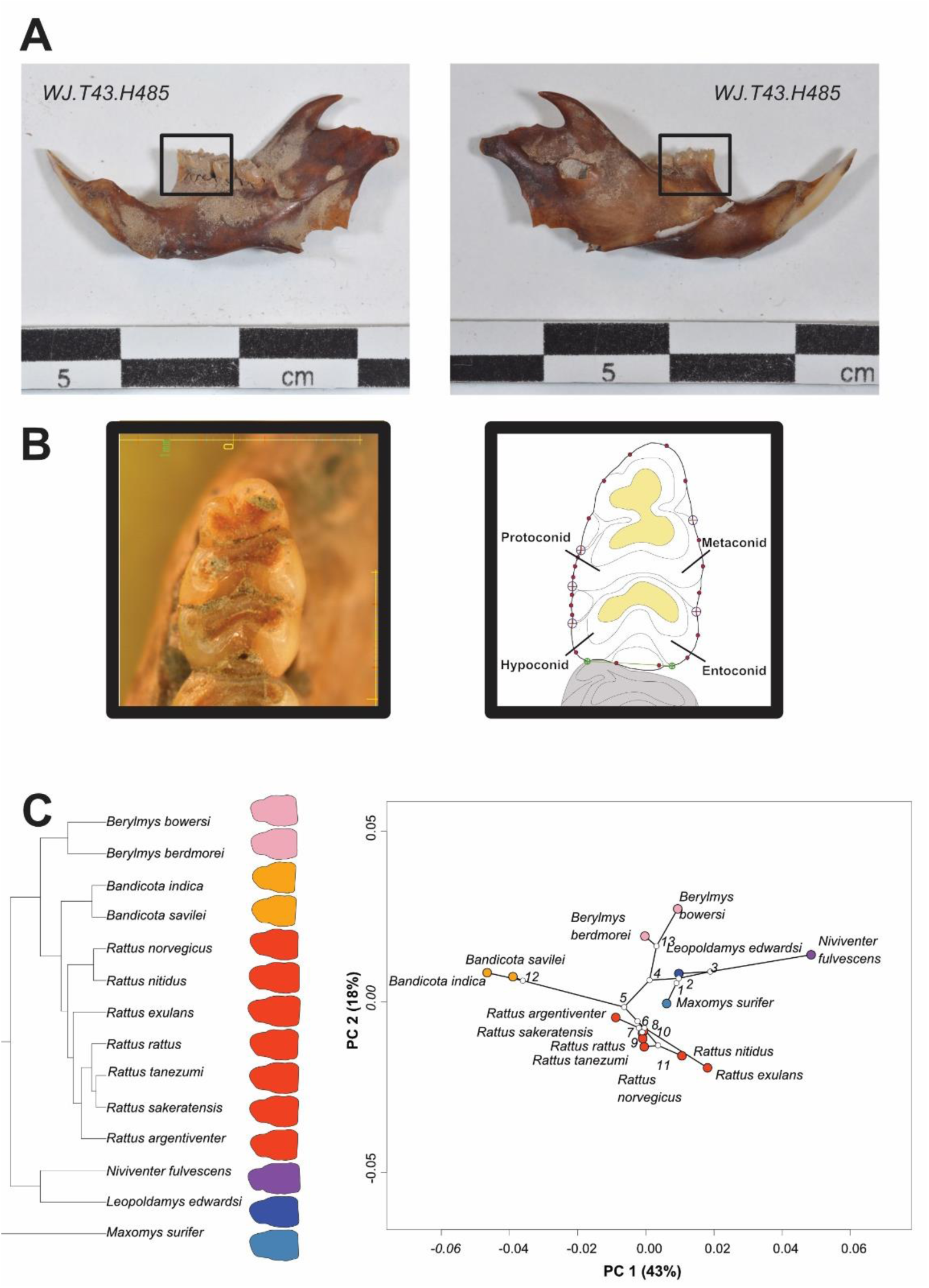
**(A)** Lingual (left) and labial (right) view of the *Rattus* sp. mandible found in the central western area of the Jiahu site in a domestic area (WJ.T43.H485). The first lower molar (m1) is framed. **(B)** Left, ortho-photography of the m1. Right, 2D GMM protocol for the m1 from Hulme-Beaman et al. 2018. **(C)** Phylogenies mapped to morphospace from Hulme-Beaman et al. (2018): Left, phylogeny of all *Rattini* species including multiple genera and the mandibular first molar’s mean shape of each species with a different colour for each genus; Right, phylogeny of all species mapped to morphospace with colours corresponding to left.

To ascertain the contemporaneity of the *Rattus* remains with the settlement deposits and exclude the possibility of contamination due to burrowing, we selected samples for direct radiocarbon AMS dating performed by Peking University. To avoid the destruction of the mandibles, we decided to provide 8 samples of *Rattus* sp. from the 4 units using appendicular bones. We obtained a direct date (lab code: BA13023) on two *Rattus* sp. femora of 6835 +/- 30 BP (95%: 5785-5642 cal BC/7735-7592 cal BP with Oxcal 4.4.4 using IntCal 2020), confirming that rat remains from Jiahu were deposited at least during the later occupation phase of the settlement.

To perform the *Rattus* species identification of the 13 archaeological mandibles, we applied 2D GMM molar shape analysis following a previously tested protocol on 395 adult and subadult specimens of the *Rattini* tribe (Hulme-Beaman et al. 2018), representing 14 species (Pagès et al. 2010) which were all genotyped and curated during the CERoPath Project (http://www.ceropath.org/) (See Hulme-Beaman et al 2018). Ortho-photographs of the first mandibular molar (m1) were taken from the 13 archaeological rats remains examined in this paper (See Figure 2B). It was not possible to use the same microscope for the archaeological and reference material, therefore we assume the biological signal is greater than between lens differences (additionally centroid size should be minimally affected by inter-equipment differences). From the m1 images we collected 5 fixed landmarks and 95 sliding semilandmarks on the 2D outline of the m1 occlusal view (see Figure 2B) following Hulme-Beaman et al. (2018) using TpsUtil and tpsDig (Goodall 1991). The size of the molar outline was quantified using centroid size (CS), which is the square root of the sum of the squared distances from the centroid of the configuration to each landmark in the configuration. The landmark data was then superimposed using a Generalised Procrustes Analysis (GPA); this was carried out to remove factors of size, position and orientation from the specimen configurations (Goodall 1991). Semilandmarks were slid using Bookstein’s minimum bending energy method (Bookstein 1997). As molar form (size+shape) rather than molar shape only was previously demonstrated to provide greater taxonomic signal among rat species (Hulme-Beaman et al. 2018), form analyses were carried out following Mitteroecker et al. (2005) for group assignment prediction of these archaeological rat specimens. This is conducted by carrying out a principal component analysis (PCA) on the combination of the log CS and rotated landmark configurations from the GPA.

The following statistical procedures were carried out on form PCs. As linear discriminant analyses require more observations (individuals per group) than variables (e.g. Mitteroecker & Bookstein 2011), the dimensionality of the data was reduced using a stepwise discriminant approach where reference datasets were resampled to equal sample size for 999 iterations. The unknown archaeological specimens were removed and the percentage correct-cross-validation (%CCV) for the remaining known balanced groups was calculated for each consecutive set of PCs up to the minimum sample size for the reference groups (13 *Rattus nitidus* and therefore 13 PCs in this study). As the reference datasets were iteratively resampled to equal sample size 999 times, the %CCV varied between iterations. Therefore, the mean %CCV values were taken at each step. The posterior probabilities of the unknown archaeological specimens to each species were calculated for each consecutive combination of PCs, and both the species with the highest posterior probability and the posterior probability for each species in each iteration were returned. The frequencies for the species with the highest probability in each iteration was plotted in a bar chart and the raw posterior probabilities for each iteration were collected and used to plot a density plot to show relative confidence in the probabilities (a single high peak can be interpreted as the posterior probability as being consistent, but a flat or multi peak distribution in posterior probabilities indicates variation in results across resampling iteration, see SI for these results). The number of consecutive PCs that achieved the maximum mean %CCV was then used for assigning species identification to the archaeological specimens but these were considered in the context of the accompanying posterior probability confidence distributions and any conflicting identifications.

All analyses were carried out in R (R core Team 2021) using functions from the base software and from the packages Morpho (Schlager, 2017) and Geomorph (Adams et al. 2024; Baken et al. 2021; Collyer & Adams 2018, 2024) and ggplot (Wickham 2016). The data, metadata and R code with additional in detailed analyses for achieving the results produced here are available in the following **data depository** (https://doi.org/10.34847/NKL.EEFD1G95).

## Results and discussion

The taxonomic identifications of the archaeological *Rattini* (Table 1, Figure 3, data depository) show that the three rats from the oldest and southernmost pre-agricultural site, Zengpiyan, were identified as a chestnut white-bellied rat (*Niviventer fulvescens)*, small white-toothed rat *(Berlylmys berdmorei)* with high confidence, and either *Maxomys surifer/ Niviventer fulvescens* with poor confidence. Of the six specimens from the early Neolithic site of Jiahu, located at an intermediate latitude, five were identified with high levels of confidence to the brown rat (*R. norvegicus)* and one specimen was identified as *Berlylmys berdmorei* with very poor confidence (40%). Of the four specimens from the northernmost site Xichengyi, two were identified as *R. norvegicus*, one as white-footed Indochinese rat (*R. nitidus)* and one as Edward’s giant rat (*Leopoldamys edwardsi)*. The other commensal rat species from this region, the Asian house rat (*Rattus tanezumi*), considered part of the same species complex as the black rat (Aplin et al. 2011), was not identified amongst these specimens.

**Figure 3:**
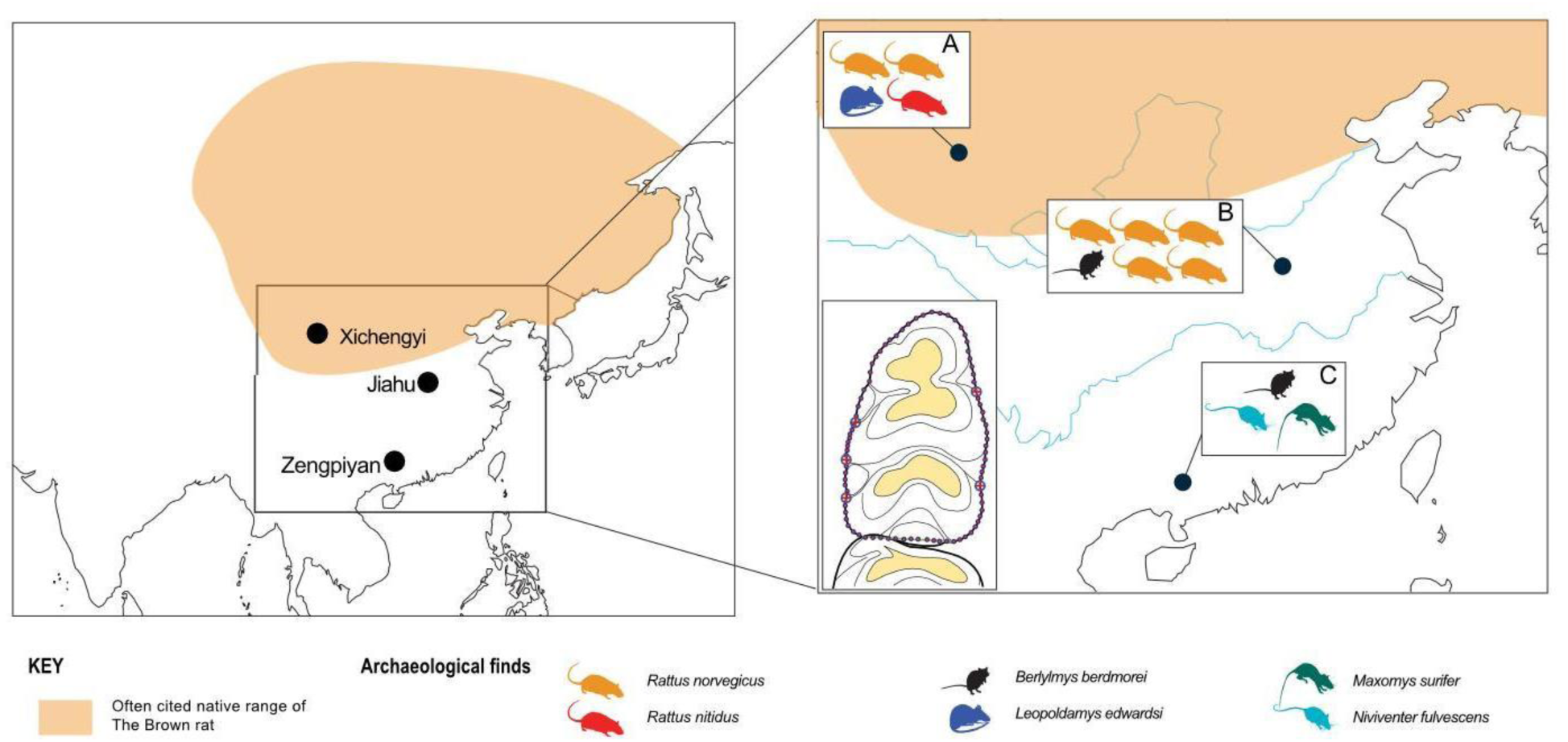
Taxonomic identification with 2D geometric morphometrics of *Rattini* dental remains (first mandibular molar) from the Chinese archaeological sites of Xichengyi (A), Jiahu (B) and Zengpiyan (C).

**Table 1:** Taxonomic identification of the thirteen archaeological mandibles from 2D GMM of mandibular first lower molar outline using a classification approach from a linear Discriminant model (see data depository https://doi.org/10.34847/NKL.EEFD1G95 for supplementary information).

| Site | Contexts | GMM ID | Unbalanced_form_95var | IDvals | OptLDA_balanced_form_13PCs | Note |
| --- | --- | --- | --- | --- | --- | --- |
| Zengpyian | 2001GGZDT4(26) | ZPY-M1-004 | *Niviventer fulvescens* | 0.996 | *Niviventer fulvescens* |  |
| Zengpyian | 2001GGZDT6(24) | ZPY-M1-005 | *Berylmys berdmorei* | 0.999 | *Berylmys berdmorei* |  |
| Zengpyian | 2001GGZDT6(19) | ZPY-M1-003 | *Maxomys surifer* | 0.52 | *Maxomys surifer*/<br>*Niviventer fulvescens* | low confidence |
| Jiahu | T43H485 | JH-M1-001 | *Rattus norvegicus* | 0.801 | *Rattus norvegicus* |  |
| Jiahu | WJT41H494 | JH-M1-002 | *Rattus norvegicus* | 0.999 | *Rattus norvegicus* |  |
| Jiahu | WBJT41H485 | JH-M1-003 | *Rattus norvegicus* | 0.875 | *Rattus norvegicus* |  |
| Jiahu | WJT41H490 | JH-M1-004 | *Rattus norvegicus* | 0.97 | *Rattus norvegicus/nitidus* | low confidence |
| Jiahu | WJT42H486 | JH-M1-005 | *Rattus norvegicus* | 0.981 | *Rattus norvegicus* |  |
| Jiahu | WJT42H486 | JH-M1-006 | *Berylmys berdmorei* | 0.405 | *Berylmys berdmorei* | low confidence |
| Xichengyi | GZH_H15-2-A852 | GZH_M1_9 | *Rattus nitidus* | 0.949 | *Rattus nitidus* |  |
| Xichengyi | GZH_H15-2-A853 | GZH_M1_10 | *Rattus norvegicus* | 0.999 | *Rattus norvegicus* |  |
| Xichengyi | GZH_H15-2-A854 | GZH_M1_11 | *Rattus norvegicus* | 0.999 | *Rattus norvegicus* |  |
| Xichengyi | GZH_T0301_4C_A355 | GZH_M1_12 | *Leopoldamys edwardsi* | 0.959 | *Leopoldamys edwardsi* | low confidence |

The *Rattini* diversity we found in Zengpiyian is the first insight into the rodent diversity of this area during the early Holocene. No specimens of the *Rattus* genus have been found but our sample is very small and it is more than likely that it does not represent the full potential diversity in this area. As an example, the more thorough excavation dedicated to paleoenvironmental studies in a 9000 years old deposit in Yunnan found six species of rodent, within only 94 NISP of small mammals, including three *Rattus* species (*R. tanezumi, R. nitidus* and *R. rattus*) as well as one species of *Niviventer* and one species of *Leopoldamys* (Jin et al. 2012). Unfortunately, we do not have access to further information regarding this material to discuss whether the rodents identified at Zengpiyan were used for food, were humans’ commensals or were brought into the deposit by raptors.

Situated in north central China, Jiahu falls close to the often-proposed native range for *R. norvegicus*, namely northern China and Mongolia (Hedrich 2006). The identification of this species at Jiahu provides the first archaeological evidence to confirm the extent of its range prior to mass human-mediated expansion. The Neolithic site of Jiahu, which was occupied for several millennia, has provided evidence of Neolithic technological advances in east Asia, including intense grain-based agriculture (Juzhong & Xiangkun 1998). Grain production at this site includes extensive evidence for rice (Juzhong & Xiangkun 1998) and there is some tentative isotopic evidence to suggest millet might be present (Hu et al. 2006). The organisation and complexity of the site, which includes hundreds of buildings and cellars and in later periods separate areas for working and residence (Zhang et al. 1999), is such that only an anthropophilic animal would have entered such an anthropogenic environment (Hulme-Beaman et al. 2016). Therefore, the occurrence of rodent taxa directly dated to c. 7700 cal BP within living areas with habitations, food storage and rubbish wastes indicates the likelihood of an emerging commensal relationship of brown rats with the early Neolithic villages in China.

This occurrence of *Rattus norvegicus* as part of Neolithic niche construction in China provides a strong argument for the commensal trajectory hypothesis of the leopard cat (*Prionailurus bengalensis*) c. 5500 cal BP, implying that the latter were attracted into early Neolithic villages by the proliferation of brown rats, themselves attracted there by the crops and their refuse (Vigne et al. 2016). Although the domestication and/or commensal status of these finds has been debated (Bar-Oz et al. 2014; Hu & Marshall 2014), the presence of these cats in anthropogenic contexts within organised and extensive agricultural settlements is considered the direct result of these cats feeding upon commensal rodents and is thought to indicate more widely the emerging commensal relationships that occur with human activity in the landscape (Cucchi et al. 2020).

The northern site of Xichengyi also provided evidence of the presence of *R. norvegicus* and this site comfortably sits within the assumed native range of this species. *R. nitidus* was also identified from here, but this site is situated further north than the modern range of *R. nitidus,* suggesting a shift of the distribution area of this species since the early Holocene. Xichengyi has indeed seen some extreme changes in its local ecology and surrounding environment, with extensive evidence for desertification leading to rapid abandonment (Wang et al. 2014). Environmental evidence indicates that during periods of occupation the site was wetter and more humid, which may have facilitated the range expansion of *R. nitidus* to the north.

### Putting these results into the global context

First, our results provide a new argument to support the parallels in evolutionary trajectories of synanthropic behaviour during the Neolithic transition between East and West Asian (Vigne 2015, 2019). In each case, crop accumulations in the earliest hamlets or villages created a new anthropogenic ecosystem where murids (brown rats in the East and house mouse, *Mus m. domesticus*, in the West; Weissbrod et al. 2017) and their specific predators, small felids (*P. bengalensis* in the East, *Felis s. lybica* in the West) were attracted, initiating a new evolutionary trajectory in which all three species – human, murid and cat – were tightly connected (Cucchi et al. 2020) in an hybrid community (Stépanoff & Vigne, 2018; Bogaard et al. 2021). However, it has been shown that Southwestern Asian mice became commensal with human sedentism rather than as a consequence of the start of agriculture (Weissbrod et al. 2017; Cucchi et al. 2020). Unfortunately, we do not have enough information regarding the contexts to determine whether the emergence of their commensalism resulted from the beginning of sedentism or cultivation.

The specimens we identify here are undoubtedly from Neolithic contexts with extensive human activity and anthropogenic change in the landscape. The site of Jiahu presents some of the earliest evidence for many Neolithic innovations in China that would have created a new ecological niche in the brown rat habitat, including an increase in consumable anthropogenic resources via grain and food storage (Juzhong & Xiangkun 1998). However, it is unlikely that commensal dependence developed immediately: given the intense level of competition observed in the commensal niche (e.g. rapid replacement of black rats by brown rats, Smith 1768) and relatively low level of human density in the overall range of the brown rat at this time, dependence was probably not established until much later (see again the example of the house mouse; Cucchi et al. 2020).

The evolutionary history of the development of commensal dependence, and the subsequent rapid increase in population size that is being estimated and tracked using genetic data, is as complex and varied as the uptake and fixation of human technological innovations. For the brown rat, Neolithic agricultural developments likely represented the first door to commensalism; urbanism alone, with its increase in human density and waste c. 4000 cal BP (Fig. 1.) may not have been enough for the second step. Subsequent intensification of urbanism, specifically with complex sewer systems, could have been the second requirement for the advancement of the fossorial brown rat into increased and regular human contact such that it could establish dependence. Yet, when brown rats reached Europe, they were often found in places without developed urbanism and complex sewer systems, suggesting that other ecological factors might have prevailed for the success of the brown rat biological invasion that need to be further understood.

## Conclusion

The brown rat has had one of the greatest impacts on human society of any animal and much of this impact stems from its prevalence due to its commensal behaviour. The relatively late arrival of the brown rat to Europe, well over a thousand years after the arrival of the black rat, gives the false impression that this species was late in adapting to the human niche and commensal behaviour. The direct evidence we present here for an abundance of brown rats deep within settlement contexts and human refuse deposits at early and intensively occupied Neolithic sites clearly demonstrates that the emergence of commensal behaviour in brown rats occurred several thousand years earlier than previously thought. This result frames the earliest adaptations of this species to the human environment. It also provides insights into the likely native range of this species, indicating that the species is either native to or expanded into central China from an early date. This then raises a range of questions regarding the timing, nature and pace of the expansion of the brown rat to the rest of the world – particularly why there was such a long apparent lag between commensalism and expansion out of eastern Asia. Further examination of both the European and East Asian archaeological record is required to understand the timing and environmental factors behind this species’ expansion and subsequent commensal dominance across much of the world.

## Acknowledgements

We would like to thank Dr Chengming Huang, Beijing Museum of Zoology, Chinese Academy of Science, Beijing; Professor Hu Yaowu, Institute of vertebrate Paleontology and Paleoanthro-pology of the Chinese Academy of Science, Beijing. We are also most grateful to Yannik Chaval and Serge Morand for giving us access to the *Rattini* skulls collected during the CERoPath project (http://www.ceropath.org/).

## Footnotes

1 Although this is the earliest evidence in the literature, we have concerns around the validity of this identification

